## Supplementary Information for "Structural mechanism of GTPase-powered ribosome-tRNA movement"

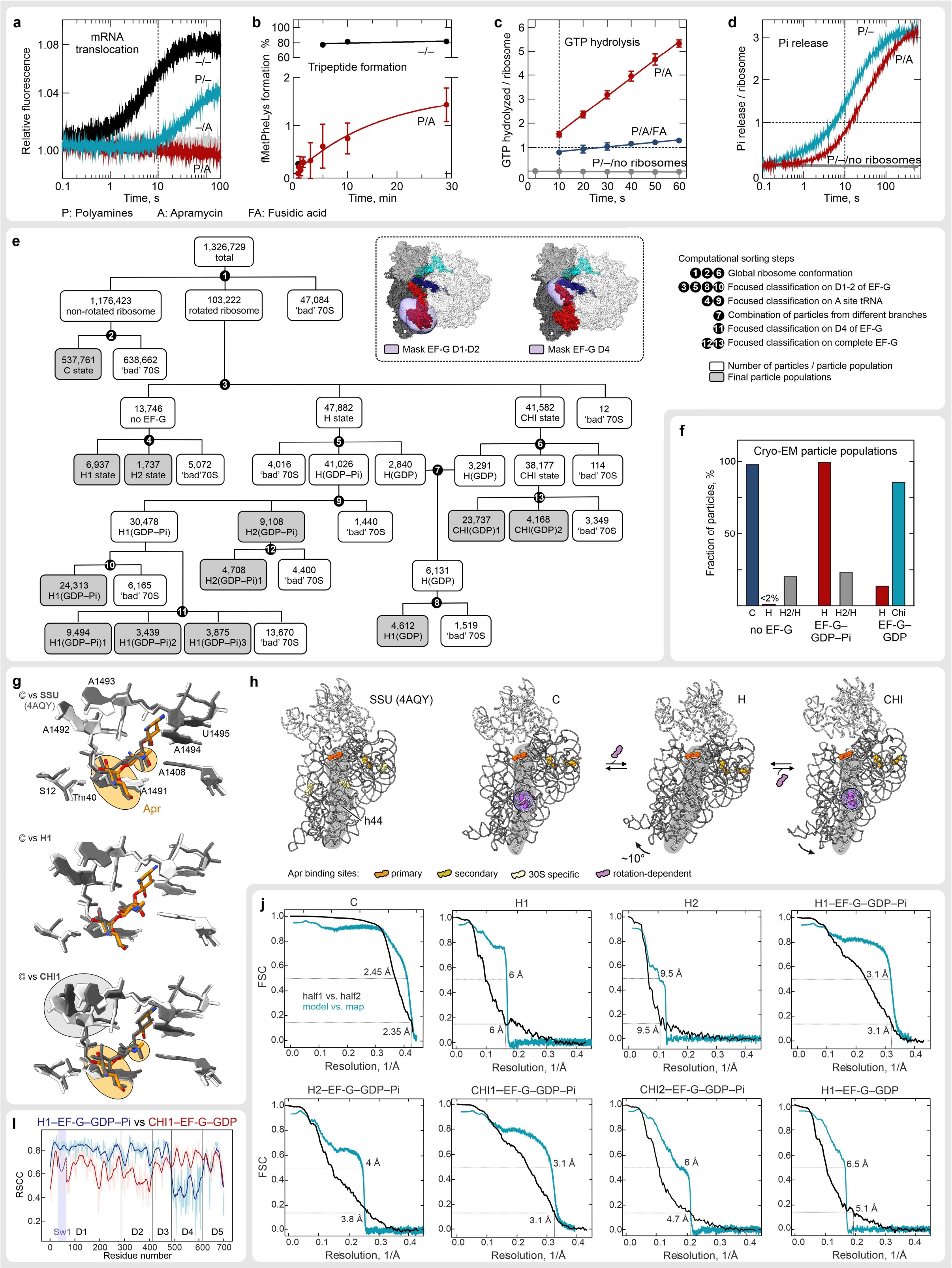


**Supplementary Fig. 1 | Biochemical characterization and cryo-EM analysis of translocation and Apr binding to the ribosome (4°C).**

**a)** Effect of polyamines (P) and Apr (A) on mRNA translocation. Translocation was monitored using a fluorescence-labeled mRNA[35]. The rate of translocation at 4°C is 0.13 $s^{-1}$ (black), in the presence of polyamines 0.016 $s^{-1}$ (cyan; 0.6 mM spermine and 0.4 mM spermidine). Addition of Apr (50 µM) abolishes translocation (gray without and red with polyamines).

- **b)** Translocation monitored by tripeptide formation. Note the split Y-axis used to visualize the low translocation activity in the presence of Apr.
- **c)** GTP hydrolysis by EF-G at multiple-turnover conditions. The close to 1:1 stoichiometry of single round GTP hydrolysis was validated by addition of the antibiotic fusidic acid, which blocks EF-G on the ribosome after one round of translocation[52].
- **d)** Pi release. Pi release was measured in the absence and presence of Apr under the same conditions as GTP hydrolysis.
- **e)** Sorting of cryo-EM data and masks used for focused classification on EF-G domains (dashed box). See Methods for details.
- **f)** Ribosome population distribution depending on the nucleotide-binding state of EF-G derived by cryo-EM particle sorting. EF-G–GDP–Pi binding alters the ribosome population from predominantly C to exclusively H, but does not significantly shift the equilibrium between the H1 and H2 states.
- **g)** Primary Apr binding site in the SSU decoding center.
  Top: Differences between Apr binding to the C state (2.35 Å resolution, this paper) and to free SSUs, as seen in the 3.5 Å crystal structure of the *Thermus thermophilus* SSU[20]. The configurations of Apr differ in the rotation of the terminal glucopyranose ring and a flip of the methylamino side group, resulting in different interaction patterns, even though the conformations of the decoding center are very similar in the two structures.
  Middle: The decoding center and Apr are similar in the C and H1 states, despite the large difference in global ribosome conformation.
  Bottom: Rearrangement of Apr between the C and CHI states. The glucopyranose ring of Apr rotates to accommodate the changes in A1492 and 1493 upon release of the mRNA-tRNA complex from the SSU body.
- **h)** Secondary Apr binding sites. Shown are Apr binding sites in the free SSU subunit (SSU) (Matt et al., 2012), state C, H1 state with EF-G–GDP–Pi, and CHI1 state with EF-G–GDP. Apr binds to the secondary binding site on h44 of 16S rRNA only at low SSU rotation degrees.
- **i)** Change in EF-G dynamics upon Pi release. The model to map real-space-cross-correlation (RSSC) is plotted for each EF-G residue of the H1–EF-G–GDP–Pi (blue) and the CHI1–EF-G–GDP (red) structures; darker lines correspond to the smoothened curves using the Savitzky–Golay filter. The more dynamic sw1 region in CHI1 state was modelled at 5 Å resolution and its RSSC determined at the same resolution.
- **j)** Fourier-Shell-Correlation (FSC) curves for final cryo-EM reconstructions.

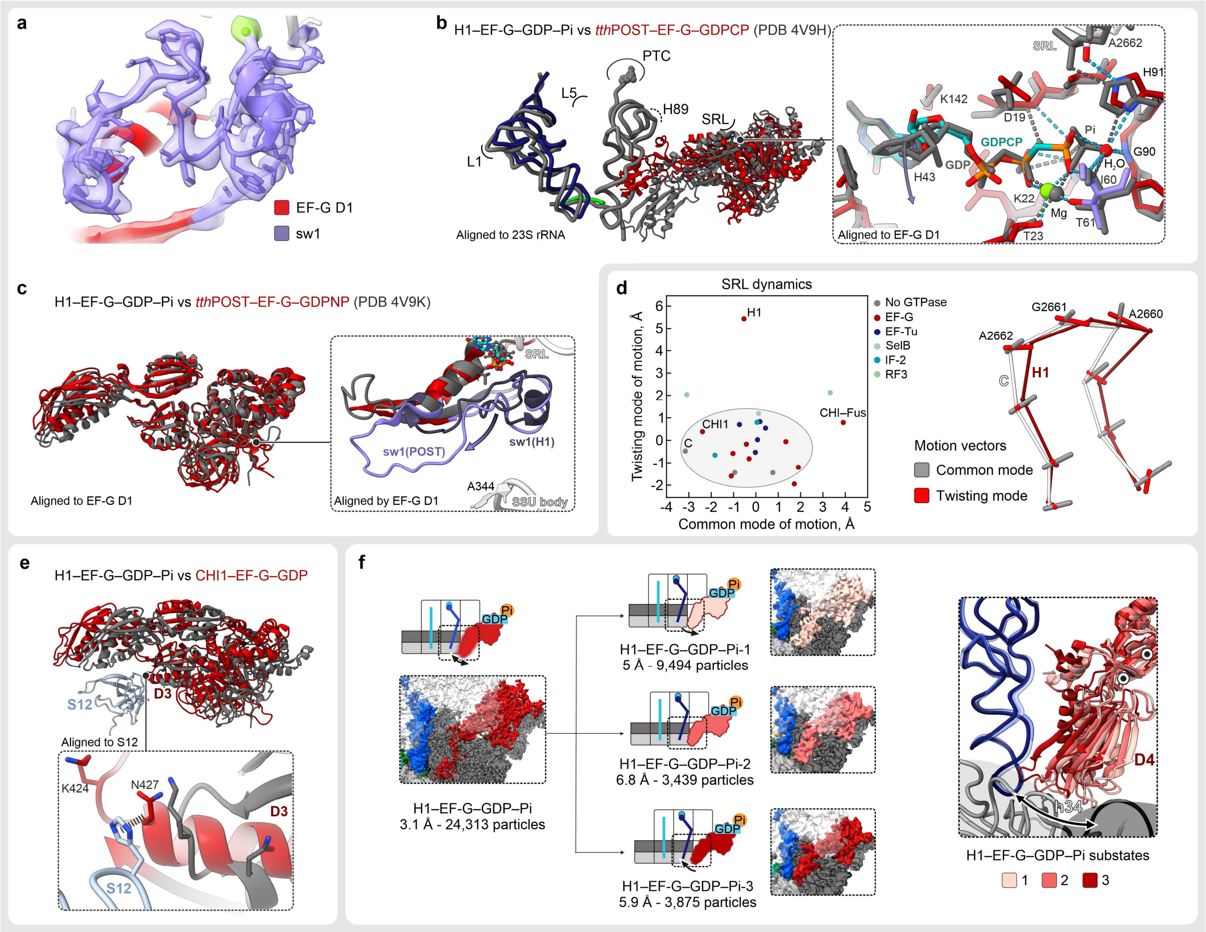


**Supplementary Fig. 2 | Details of the H–EF-G–GDP–Pi structures.**

**a)** Cryo-EM density for the sw1 region in the H1–EF-G–GDP–Pi structure at 3.1 Å.

**b)** Comparison of the H1–EF-G–GDP–Pi structure with the structure of EF-G–GDPCP bound to the POST ribosome[15] (PDB 4V9H). The global EF-G position on the ribosome, including the SRL, is different (left), but the overall EF-G conformation is similar (RMSD 2.0 Å with sub-state 3 of H1–EF-G–GDP–Pi excluding sw1). Also, the nucleotide binding pocket appears remarkably similar in the two structures (right). The sw1 region is mostly disordered in EF-G–GDPCP, except for the short helix (residues 52-64) that is structured.

**c)** Structural comparison of H1–EF-G–GDP–Pi and POST–EF-G–GDPNP [13] (PDB 4V9K). The overall conformation of EF-G is similar (left), despite the different global position on the ribosome (not shown; similar in refs.[13] and[15]). The sw1 region (right) adopts an extended conformation in the POST complex and points away from the SSU.

**d)** Unique conformational change of the SRL in EF-G–GDP–Pi. Left: Structural dynamics of the SRL. The two major conformational modes of the SRL were determined by principal component analysis of ribosome structures with and without translational GTPases (Methods). The first "common" mode describes the prevalent dynamics of the SRL, the second "twisting" mode reflects the unique change observed in the H–EF-G–GDP–Pi state. Right: Conformational dynamics of SRL residues (solid bars) projected onto the SRL backbone structure in the H1–EF-G–GDP–Pi (dark red) and C state (white). The common mode (grey bars) uniformly affects the entire SRL structure, whereas the twisting mode (light red bars) mostly affects the crucial bases A2660-A2662.

**e)** Dynamic interactions of EF-G D3 with protein S12. Upon Pi release EF-G slides on S12 switching the contact with His76 of S12 from Lys424 in the H1 state to Asp427 in the CHI1 state.

**f)** Conformations of EF-G–GDP–Pi D4 in H1. Left: The dynamics of D4 in the H1 state were resolved by sorting the corresponding cryo-EM data into three sub-states and computing separate cryo-EM maps. Right: The superposition shows the three distinct sub-states of D4. Notably, the broad conformational range sampled by D4 has no effect on the ribosome conformation, or tRNA positions. D4 moves largely independent of the other EF-G domains, facilitated by flexible hinge regions (black dots).

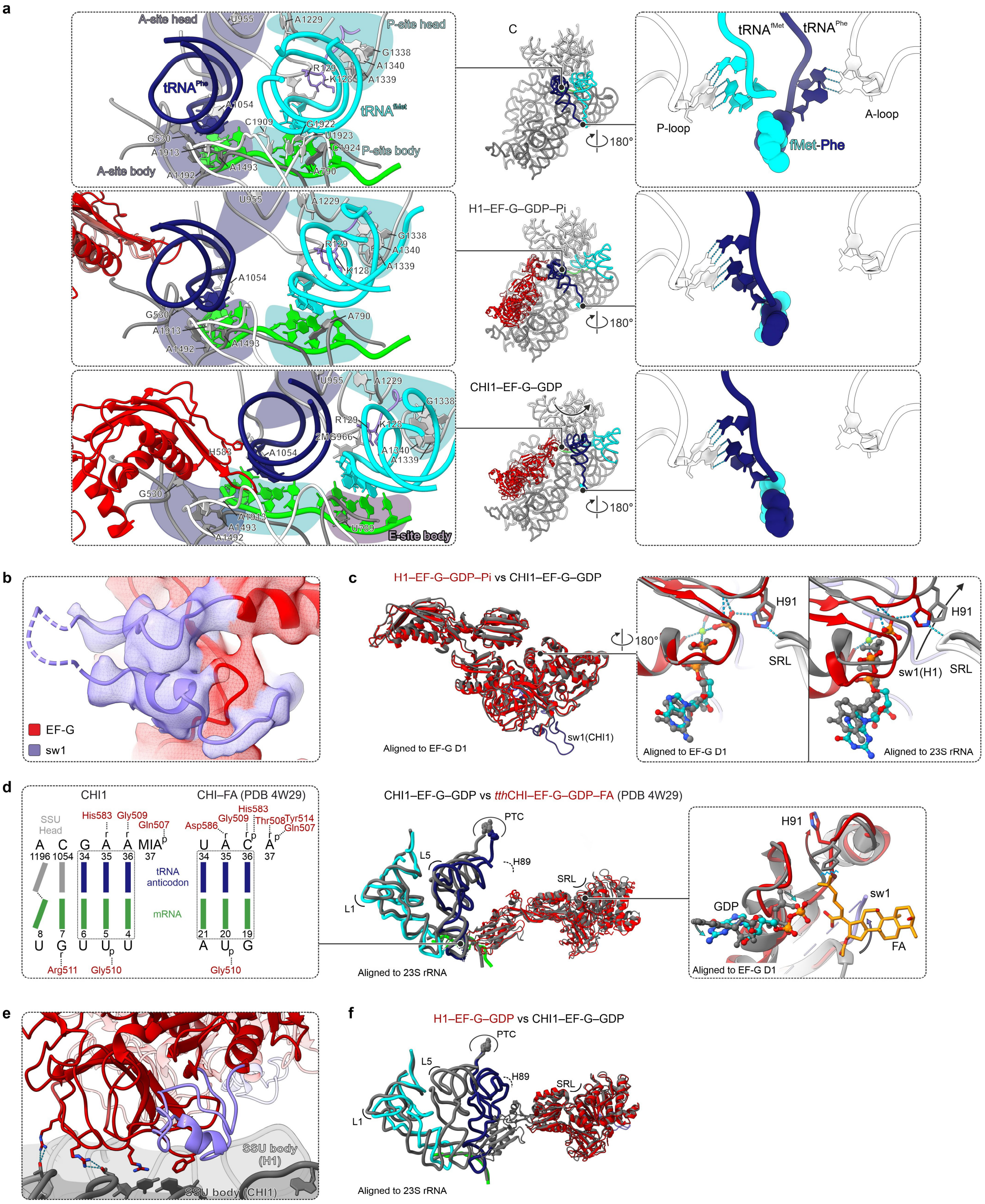


**Supplementary Fig. 3 | Details of the CHI1–EF-G–GDP structure.**

**a)** Movement of the tRNAs from the C (top) to H1 (middle) to CHI1 (bottom) state. Close-ups of the decoding center (left) and the peptidyl transferase center (right). On the SSU, the A-site fMetPhe-tRNA$^{Phe}$ moves from the A site in C and H1 states to the ap state in CHI1, while the P-site tRNA$^{fMet}$ moves from the P site to the pe state. On the LSU, fMetPhe-CCA is in the A site in C and in the P site in the H1 and CHI1 states.

**b)** Cryo-EM density of the refolded sw1 region in the CHI1–EF-G–GDP state. By low-pass filtering to 5 Å we were able to model the backbone of nearly the entire sw1, except the dashed region. The short α-helix (residues 52-57) in sw1 remains unchanged after the large-scale sw1 refolding upon Pi release.

**c)** Superposition of EF-G in H1–EF-G–GDP–Pi and CHI1–EF-G–GDP. Left: Overall EF-G conformation. Note the similarity also in EF-G D4, which is stably bound to the decoding center in CHI states, but is dynamic in H states where it samples several conformations, including that found in CHI state (the most similar sub-state 3 of H1 is shown). Right: Close-ups of Pi coordination and SRL contacts. In contrast to sw1, the conformation of sw2 (containing the catalytic His91) changes only slightly upon Pi release, whereas the contacts with the SRL change substantially.

**d)** Comparison of the CHI1–EF-G–GDP structure (3.1 Å, this paper) and the 3.8 Å crystal structure of a *T. thermophilus* CHI–EF-G–GDP complex trapped with fusidic acid [14] (PDB 4W29). Left: Contacts of the mRNA-tRNA complex in the ap site of the SSU. Center: Global positions of EF-G and tRNAs. Note the major difference in the acceptor stem position of the A-site tRNAs, which might result from the different A-site tRNAs used, fMetPhe-$tRNA^{Phe}$ in our structure vs. the peptidyl-tRNA analogue N-acetyl-Val-$tRNA^{Val}$ in the crystal structure. Right: Close up of the EF-G nucleotide binding pocket showing significant differences in the GDP position, as well as in EF-G sw1 and sw2, which may result from the presence of fusidic acid (FA) in the crystal structure.

**e)** Repositioning of EF-G D2 on the SSU shoulder upon Pi release. The superposition of CHI1–EF-G–GDP and H1–EF-G–GDP–Pi (semi-transparent in the background) using 23S rRNA illustrates the large-scale change in the position of D2 and the SSU shoulder due to back-rotation of the SSU body domain.

**f)** Pi release uncoupled from translocation. In a minor population of H1–EF-G–GDP complexes, the sw1 region is unstructured and EF-G moves in a similar way as in the CHI1 state, indicating that Pi has been released. However, the ribosome remains in the rotated state with tRNAs in hybrid states (H1). There is no density for D4, most likely due to high flexibility, which corroborates the importance of D4 progression into the decoding center to promote mRNA-tRNA movement.

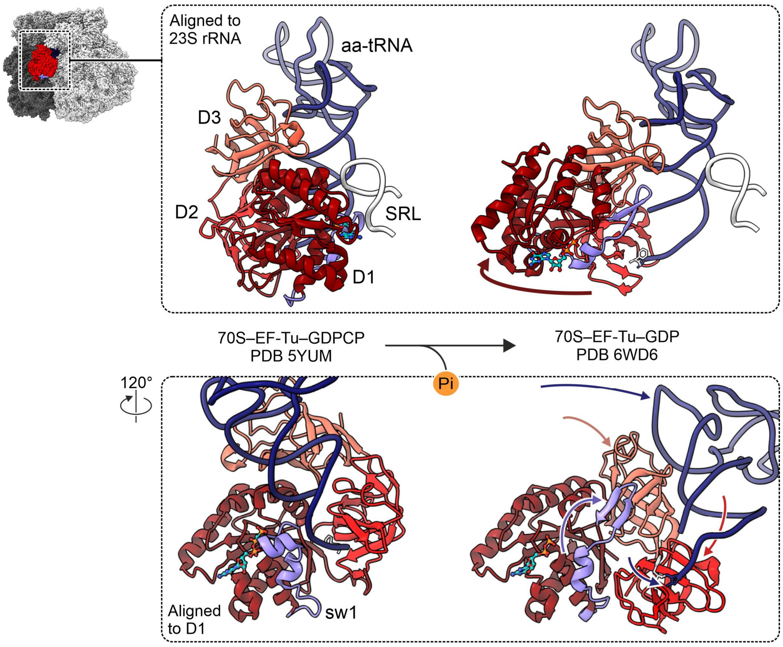


**Supplementary Fig. 4 | GTPase-driven changes of EF-Tu on the ribosome**.

Structures of EF-Tu–GDPCP in the GTPase activated state[53] (left) and EF-Tu–GDP on the ribosome[29] (right) are shown from the LSU solvent site (top) and tRNA binding site (bottom). Note the refolding of residues 55-64 of the compact sw1 into a β-hairpin upon Pi release and the substantial changes in the interdomain arrangement, which relax interactions of EF-Tu with the aminoacyl-tRNA (aa-tRNA) and ribosome. For visual clarity, the SRL is not shown in the bottom panels.

### **Supplementary Table 1.** Cryo-EM structure determination – part 1

| **Ribosomal complex** | Classic | H1 | H2 | H1–EF-G–GDP–Pi |
|---|---|---|---|---|
| Ribosomal state | Classic tRNA state | Hybrid tRNA state 1 | Hybrid tRNA state 2 | Hybrid tRNA state 1 with EF-G–GDP–Pi |
| **Database entries** | | | | |
| EMDB ID | **YYYY** | **YYYY** | **YYYY** | **YYYY** |
| PDB ID | **XXXX** | **XXXX** | **XXXX** | **XXXX** |
| Final resolution (Å)*,◊ | 2.35 | 6 | 9.5 | 3.1 |
| **Data collection** | | | | |
| Microscope | Titan Krios | Titan Krios | Titan Krios | Titan Krios |
| Camera | Falcon III | Falcon III | Falcon III | Falcon III |
| Magnification | 59,000 | 59,000 | 59,000 | 59,000 |
| Voltage (kV) | 300 | 300 | 300 | 300 |
| Electron dose ($e^{-}$/Å$^2$) | 30 | 30 | 30 | 30 |
| Defocus range (µm) | 0.2-1.5 | 0.2-1.5 | 0.2-1.5 | 0.2-1.5 |
| Pixel size (Å) | 1.16 | 1.16 | 1.16 | 1.16 |
| **Cryo-EM reconstruction** | | | | |
| Final particles (no.) | 537,761 | 6,937 | 1,737 | 24,313 |
| Point group symmetry | *C1* | *C1* | *C1* | *C1* |
| FSC-threshold | 0.143 | 0.143 | 0.143 | 0.143 |
| Resolution (Å) | 2.35 | 6 | 9.5 | 3.1 |
| Resolution metric | gold standard FSC | gold standard FSC | gold standard FSC | gold standard FSC |
| **Atomic model refinement** | | | | |
| Resolution§ (Å) | 2.45 | 6 | 9.5 | 3.1 |
| Cumulative RSCC (%) >0.8/>0.6/>0.4 | 0.89/0.96/0.98 | 0.89/0.97/0.99 | 0.3/0.92/0.99 | 0.62/0.92/0.97 |
| Initial models used | 6YSS (70S,Phe-tRNA) | 6YSS (70S,Phe-tRNA) | 6YSS (70S,Phe-tRNA) | 6YSS (70S,Phe-tRNA) |
| | 4AQY(Apr) | 4AQY(Apr) | 4AQY(Apr) | 4AQY(Apr) |
| | 5LZD (fMet-tRNA) | 5LZD (fMet-tRNA) | 5LZD (fMet-tRNA) | 5LZD (fMet-tRNA) |
| | | | | 3J9Z(EF-G) |
| Molprobity score | 2.54 | 2.99 | 2.36 | 2.49 |
| Clashscore | 12.69 | 63.51 | 17.97 | 16.20 |
| **No. Atoms/No. Residues/RSCC** | | | | |
| Total | 147,741/11,065/0.87 | 147,241/11,075/0.86 | 147,241/11,075/0.74 | 153,165/10,762/0.79 |
| Protein | 45,743/5,954/0.86 | 45,742/5,954/0.84 | 45,742/5,954/0.70 | 51,187/6,657/0.77 |
| Nucleic | 101,460/4,727/0.87 | 101,460/4,727/0.88 | 101,460/4,727/0.79 | 101,461/4,727/0.81 |
| EF-G | - | - | - | 5,444/703/0.72 |
| **B-factors** | | | | |
| Protein | 79.51 | 64.92 | 44.64 | 47.47 |
| Nucleotide | 72.70 | 69.59 | 47.80 | 49.65 |
| Ligands, Ions | 72.79 | 45.53 | 30.97 | 44.01 |
| **R.m.s. deviations** | | | | |
| Bond lengths (Å) | 0.011 | 0.009 | 0.008 | 0.009 |
| Bond angles (°) | 1.061 | 1.069 | 0.904 | 0.969 |
| **Ramachandran plot** | | | | |
| Favored (%) | 91.90 | 81.52 | 87.71 | 88.54 |
| Allowed (%) | 7.45 | 18.32 | 12.10 | 11.16 |
| Disallowed (%) | 0.65 | 0.15 | 0.19 | 0.31 |

*For model refinement, maps at ≤3.1Å resolution were resampled to 512×512×512 pixels, corresponding to a pixel size of 0.6525Å
◊Resolution used for atomic model refinement and interpretation.
§Resolution at which map-model FSC=0.5.

### **Supplementary Table 2.** Cryo-EM structure determination – part 2

| **Ribosomal complex** | H2–EF-G–GDP–Pi | CHI1–EF-G–GDP | CHI2–EF-G–GDP | H1–EF-G–GDP |
|---|---|---|---|---|
| Ribosomal state | Hybrid tRNA state 2 with EF-G–GDP–Pi | Chimeric tRNA state 1 with EF-G–GDP | Chimeric tRNA state 2 with EF-G–GDP | Hybrid tRNA state 1 with EF-G–GDP |
| **Database entries** | | | | |
| EMDB ID | **YYYY** | **YYYY** | **YYYY** | **YYYY** |
| PDB ID | **XXXX** | **XXXX** | **XXXX** | **XXXX** |
| Final resolution (Å) *,◊ | 4 | 3.1 | 6 | 6.5 |
| **Data collection** | | | | |
| Microscope | Titan Krios | Titan Krios | Titan Krios | Titan Krios |
| Camera | Falcon III | Falcon III | Falcon III | Falcon III |
| Magnification | 59,000 | 59,000 | 59,000 | 59,000 |
| Voltage (kV) | 300 | 300 | 300 | 300 |
| Electron dose ($e^{-}/Å^2$) | 30 | 30 | 30 | 30 |
| Defocus range (μm) | 0.2-1.5 | 0.2-1.5 | 0.2-1.5 | 0.2-1.5 |
| Pixel size (Å) | 1.16 | 1.16 | 1.16 | 1.16 |
| **Cryo-EM reconstruction** | | | | |
| Final particles (no.) | 9,108 | 23,737 | 4,168 | 4,612 |
| Point group symmetry | *C1* | *C1* | *C1* | *C1* |
| FSC-threshold | 0.143 | 0.143 | 0.143 | 0.143 |
| Resolution (Å) | 3.8 | 3.1 | 4.7 | 5.1 |
| Resolution metric | gold standard FSC | gold standard FSC | gold standard FSC | gold standard FSC |
| **Atomic model refinement** | | | | |
| Resolution§ (Å) | 4 | 3.1 | 6 | 6.5 |
| Cumulative RSCC (%) >0.8/>0.6/>0.4 | 0.14/0.87/0.95 | 0.4/0.86/0.96 | 0.42/0.93/0.98 | 0.4/0.86/0.95 |
| Initial models used | 6YSS (70S, Phe-tRNA) | 6YSS (70S, Phe-tRNA) | 6YSS (70S, Phe-tRNA) | 6YSS (70S, Phe-tRNA) |
| | 4AQY(Apr) | 4AQY(Apr) | 4AQY(Apr) | 4AQY(Apr) |
| | 5LZD (fMet-tRNA) | 5LZD (fMet-tRNA) | 5LZD (fMet-tRNA) | 5LZD (fMet-tRNA) |
| | 3J9Z(EF-G) | 3J9Z(EF-G) | 3J9Z(EF-G) | 3J9Z(EF-G) |
| Molprobity score | 2.54 | 2.99 | 2.36 | 2.49 |
| Clashscore | 23.55 | 15.98 | 91.40 | 16.81 |
| **No. Atoms/No. Residues/RSCC** | | | | |
| Total | 152,719/11,387/0.87 | 153,082/11,703/0.86 | 152,692/11,387/0.74 | 151,321/11,205/0.79 |
| Protein | 51,178/6,656/0.86 | 51,148/6,651/0.84 | 51,188/6,657/0.70 | 49,795/6,476/0.77 |
| Nucleic | 101,462/4,727/0.87 | 101,437/4,726/0.88 | 101,437/4,726/0.79 | 101,461/4,727/0.81 |
| EF-G | 5,444/703/0.57 | 5,444/703/0.64 | 5,444/703/0.62 | 4,052/522/0.44 |
| **B-factors** | | | | |
| Protein | 37.16 | 39.83 | 272.16 | 46.73 |
| Nucleotide | 45.37 | 45.22 | 243.59 | 49.65 |
| Ligands, Ions | 34.42 | 51.32 | 289.46 | 32.49 |
| **R.m.s. deviations** | | | | |
| Bond lengths (Å) | 0.007 | 0.007 | 0.015 | 0.007 |
| Bond angles (°) | 0.998 | 0.922 | 1.760 | 0.887 |
| **Ramachandran plot** | | | | |
| Favored (%) | 86.23 | 90.09 | 90.11 | 88.25 |
| Allowed (%) | 13.65 | 9.50 | 9.63 | 11.53 |
| Disallowed (%) | 0.12 | 0.41 | 0.26 | 0.22 |

*For model refinement, maps at ≤3.1Å resolution were resampled to 512×512×512 pixels, corresponding to a pixel size of 0.6525Å
◊Resolution used for atomic model refinement and interpretation.
§Resolution at which map-model FSC=0.5.

**Supplementary Movie 1 | GTPpase-powered tRNA translocation step visualized by cryo-EM.**

The movie illustrates how GTP hydrolysis by the translational GTPase EF-G results in large-scale molecular movement driving forward tRNA movement on the ribosome. The animation is based on the experimental structures; transitions between the structures are rendered by morphing.
